# Early season soil microbiome best predicts wheat grain quality

**DOI:** 10.1101/2022.09.29.510160

**Authors:** Numan Ibne Asad, Xiao-Bo Wang, Jessica Dozois, Hamed Azarbad, Philippe Constant, Etienne Yergeau

**Affiliations:** Institut national de la recherche scientifique, Centre Armand-Frappier Santé Biotechnologie, Laval, QC H7V 1B7, Canada; State Key Laboratory of Grassland Agroecosystems, Center for Grassland Microbiome and College of Pastoral, Agriculture Science and Technology, Lanzhou University, Lanzhou 730020, People’s Republic of China; Philipps-University Marburg, Department of Biology, Evolutionary Ecology of Plants, Marburg, Germany

**Keywords:** wheat microbiome, LASSO regression, grain quality, amplicon sequencing, nitrogen cycle, community level physiological profiling

## Abstract

Previous studies have shown that it is possible to accurately predict wheat grain quality and yields using microbial indicators. However, it is uncertain what the best timing for sampling is. For optimal usefulness of this modeling approach, microbial indicators from samples taken early in the season should have the best predictive power. Here, we sampled a field every two weeks across a single growing season and measured a wide array of microbial parameters (amplicon sequencing, abundance of N-cycle related functional genes, and microbial carbon usage) to find the moment when the microbial predictive power for wheat grain baking quality is highest. We found that the highest predictive power for wheat grain quality was for microbial data derived from samples taken early in the season (May–June) which coincides roughly with the seedling and tillering growth stages, that are important for wheat N nutrition. Our models based on LASSO regression also highlighted a set of microbial parameters highly coherent with our previous surveys, including alpha- and beta-diversity indices and N-cycle genes. Taken together, our results suggest that measuring microbial parameters early in the wheat growing season could help farmers better predict wheat grain quality.

## Introduction

As the world population climbs towards the 9-billion-mark, agricultural production is under the pressure of climate change. The productivity gains from the green revolution have plateaued, traditional breeding efforts can hardly keep up, and the level of pesticide and inorganic fertilizer use is unsustainable. New solutions are needed. Integrated microbiocentric approaches to optimize plant production are promising and have often been proposed to solve some of the many problems agricultural production faces (Figuerola *et al*. 2012; Schloter *et al*.2018). Soil microorganisms play a key role in many ecosystem processes that are central to agricultural production. For instance, soil microorganisms recycle organic matter, cycle nutrients, abate abiotic stresses, change soil structure and porosity, and promote plant growth (Ortiz & Sansinenea 2022). However, although it is theoretically known how to modify microbial communities (Agoussar & Yergeau 2021), it is in practice still a very daunting task because of the complexity of the communities and their interactions. A first step towards this goal would be to create microbial-based models predicting agricultural processes, to identify clear targets and key functions or taxa to manipulate.

However, soil microbial communities are very dynamic, which makes it difficult to predict process rates and to identify key players that would be amenable to manipulation. Soil microbial communities are strongly influenced by biotic and abiotic factors, such as temperature, precipitations, and plant growth stage, which all vary in time, often in an unpredictable manner. We recently showed that dry-rewetting cycles lead to a complete overhaul of the soil microbial communities, much more than small decreases in soil water content (Wang *et al*. 2022.). Soybean and wheat growth stages were shown to profoundly influence the microbial diversity associated with the plant, often in interaction with plant compartment, plant genotype, soil water content and soil history (Moroenyane *et al*. 2021; Azarbad *et al*. 2022; Azarbad *et al*. 2002). Similarly, the effect of the genotype on root and rhizosphere microbial communities varied over time (years) and with wheat growth stages (Quiza *et al*. 2022). These microbial shifts related to plant growth stages were previously linked to changes in the composition and concentration of plant root exudates during development (Chaparro *et al*. 2013). The timing of sampling is thus expected to influence the predictive power microbial parameters, but it is still uncertain what the best sampling time would be and whether robust time-independent indicators could be identified.

Recent microbial-based modeling from our group showed that early sampling of wheat field soil microbial communities, around seeding or emergence could accurately predict wheat yield and grain baking quality obtained at the end of the growing season (Asad *et al*. 2021; Yergeau *et al*. 2020). For instance, with as little as 5 predictors, such as the abundance of archaeal ammonia-oxidizers, measured shortly after seeding in May, we were able to predict wheat grain quality with an accuracy of up to 81% (Yergeau *et al*. 2020). In contrast, different ammonium nitrate fertilization regimes did not significantly influence yields or grain baking quality. In another study encompassing 80 fields across a transect of 500km, microbial indicators from samples taken in May-June could robustly predict the wheat grain quality and yields at the end of the growing season (Asad *et al*. 2021). In line with this, earlier work showed that the growth of willows after 100 days in highly contaminated soil could be predicted by the initial soil microbial diversity (Yergeau *et al*., 2015), whereas willows Zn accumulation after 16 months of growth could be predicted by the relative abundance of specific fungal taxa present at 4 months (Bell *et al*. 2015). Therefore, it seems that the early soil microbial data can accurately predict ecosystem processes, such as plant productivity and produce quality. However, these studies did not compare microbial data taken at different timepoints, so it is unclear if early sampling has the highest predictive power in microbial-based models.

Here, we sampled the same experimental field every two weeks over the course of a single growing season. We sequenced the 16S rRNA gene and the ITS region, quantified the abundance of key N-cycle genes, and measured the community level physiological profiles as microbial indicators and linked them to grain baking quality using statistical learning approaches. Our goals were to 1) identify the most appropriate sampling date for modelling, and 2) identify robust microbial indicators linked to grain baking quality.

## Methods

### Experimental design and sampling

Four rainfall manipulation treatments were set-up in 2016 at the Armand-Frappier Sante Biotechnologie Centre (Laval, Québec, Canada) using 2m x 2m rain-out shelters that excluded passively 0%, 25%, 50%, and 75% of the natural precipitation. The rainfall exclusion treatments were performed using rain-out shelters, which were covered with various amount of transparent plastic sheeting. The rain was intercepted by the plastic sheeting and guided in a gutter and downspout and collected in 20L buckets that were manually emptied following significant rainfall events. Two wheat genotypes were seeded under these shelters (drought sensitive, *Triticum aestivum* cv. AC Nass and drought tolerant, *Triticum aestivum cv. AC Barrie*), and the experiment was replicated over 6 fully randomized blocks, resulting in 48 plots (4 treatments x 2 genotypes x 6 blocks). Seeds harvested from each of the plots were re-seeded in the exact same plot the following year. Soil was sampled every 2 weeks on May 10^th^ (seeding time, T = 0), May 24^th^, June 7^th^, June 21^st^, July 5^th,^ July 19^th^, and August 1^st^ 2018. A composite soil sample was prepared by collecting 10-cm deep soil cores from the 4 corners and the centre of each plot (4 treatments x 6 blocks x 2 cultivars x 7 sampling dates = total 336 samples). From 2016 to 2018, the average daily rainfall recorded on this site was 2.2 mm-3.5 mm. Soil water content within rainfall exclusion treatments showed significant differences among soil sampling dates (Wang *et al*. 2022).

### Amplicon sequencing and data analysis

Total genomic DNA was extracted from the 336 soil samples with the DNeasy PowerLyzer Power Soil Kit (Qiagen) following the manufacturer’s instructions. The concentration and the quality of the DNA was checked using a Nano Drop ND-1000 Spectrophotometer (Nano Drop Technologies Inc., Thermo Scientific, U.S.A.). The amplicon sequencing libraries for the bacteria and archaeal 16S rRNA gene and ITS regions were prepared according to the previously described protocols (Asad *et al*. 2021). The primers pairs used for the amplification were 515F/806R (Caporaso *et al*. 2012) and ITS1F/58A2R (Martin & Rygiewicz, 2005), for the bacterial and archaeal 16S rRNA gene and the fungal ITS region, respectively. PCR amplifications were conducted in a T100™ Thermal Cycler (Bio-Rad, U.S.A.) as previously described (Wang *et al*. 2022). PCR products were confirmed through visualization in 1% agarose gel and purified using AMPure XP beads (Beckman Coulter, Indianapolis, U.S.A.). PCR libraries were pooled together and sent to the Centre d’expertise et de services Genome Québec (Montréal, Canada) for Illumina MiSeq 2 × 250 bp amplicon sequencing as detailed previously (Wang *et al*. 2022). A total of 17,084,986 16S rRNA gene reads and 22,411,001 ITS region reads were produced. The raw sequencing data and its meta data were deposited in the NCBI BioProject under accession PRJNA686206.

Sequence pre-processing, including filtering and quality testing, was performed using UCHIME (Edgar *et al*. 2011), following previously published bioinformatic pipelines (Wang *et al*. 2022). The classification of Operational Taxonomic Units (OTUs) was performed using the RDP 16S rRNA Reference Database (Wang *et al*. 2007) and the UNITE ITS Reference Database (Nilsson *et al*. 2019). The uniformity of the amplicon sequences belonging to the same operational taxonomic units (OTUs) was tested using UPARSE (Edgar *et al*. 2013). Sample rarefaction was performed using an in-house galaxy pipeline as previously discussed (Wang *et al*. 2022.). Alpha (e.g., Shannon, Simpson, Chao1, Abundance-based Coverage Estimators), beta (Bray-Curtis dissimilarity) and phylogenetic diversity were calculated as detailed in Wang *et al* (2022).

### Quantitative real-time PCR (qPCR) and community level physiological profiling (CLPP)

We measured the abundance of the 16S rRNA gene, the ITS region, and N-cycle related genes (bacterial and archaeal *amoA, nirK, nosZ*) for the 336 samples using real-time PCR SYBR Green assays, as previously described (Asad *et al*. 2021). The Fungal:Bacterial (F:B) ratio was then calculated by dividing the ITS region abundance by the 16S rRNA gene abundance. EcoPlates colorimetric assays (Biolog, Hayward, CA) were used to measure microbial carbon use patterns with diluted soil (1/10 in water) and a 168-hour incubation, as previously described (Asad *et al*. 2021).

### Wheat grain and flour quality

Wheat grain was harvested from the 48 plots at the end of the growing season (8^th^ August 2018) and the grain and flour baking quality were analyzed in the quality control laboratory of Les Moulins de Soulanges (St-Polycarpe, QC). Four main quality indicators were used in our modeling efforts: grain protein content, grain gluten content, flour peak maximum time (PMT; time for the dough to reach its maximum consistency following hydration), flour maximum recorded torque (BEM, maximal consistency as measured as resistance to mechanical mixing) (Freund and Kim 2006). A good quality grain for bread is expected to have a high protein and gluten content. A good quality flour will have a high maximum torque (high consistency) and a short PMT (rapid to reach maximal consistency) when hydrated.

### Statistical analysis

All the statistical analyses were performed in R (v.4.1.2). To visualise the differences in the microbial community (amplicon and CLPP datasets) across sampling dates, treatments, and cultivars, we used the function *cmdscale* of the vegan package to produce principal coordinate analysis (PCoA) based on the Bray-Curtis dissimilarity index. The effect of sampling date, treatments, block, genotypes on the microbial community structure and carbon utilisation patterns was tested using permutational multivariate analysis of variance (PERMANOVA) based on the Bray-Curtis dissimilarity index (*adonis2* function of the vegan package). Three-way repeated measures analysis of variance (rmANOVA) using the *aov* function was used to test for significant differences in alpha diversity, N-cycle related genes and ITS region and 16S rRNA gene abundance. The normality of the residuals was examined graphically using *ggqplot* and was tested by the Shapiro-Wilk test using the *shapiro*.*test* function. If the data did not meet the requirements of the tests, it was log or square root transformed. The homoscedasticity of the data was evaluated using the Mauchly’s sphericity test of the *rstatix* package. Correlation analyses between microbial parameters and wheat grain quality were performed with the *cor*.*test* function together with the *p*.*adjust* function to adjust the p-value with the Benjamin-Hochberg correction for multiple tests.

### Predictive modeling

Our goal was to model grain quality (protein, gluten, BEM and PMT) using the microbial indicators measured (bacterial and fungal alpha diversity, bacterial and fungal beta-diversity, carbon utilization patterns, F:B ratio, and N-cycle gene abundance), for each sampling date separately to find the optimal sampling date for modeling. Since our PERMANOVAs revealed that the two wheat genotypes harbored significantly different microbial communities, we modeled them separately. This resulted in 14 different microbial datasets containing each 24 samples. We excluded outlier data points using the *rstatix* package.

To reduce the dimensionality of the 16S rRNA gene and ITS region amplicon OTU tables and of the microbial carbon usage, we performed a procedure called orthogonalization. In brief, we performed a principal component analysis (*PCA* function of the FactomineR package) on Hellinger-transformed (*decostand* function of vegan package) OTU tables or carbon usage patterns and used the 5 first principal components in the models. Individual OTUs and carbon substrates were then correlated to these 5 components to have an idea of the taxonomic composition of the OTUs or carbon substrates influencing each of the components. We kept OTUs and carbon substrates with correlation having a P<0.05. For the OTUs, a taxonomic summary at the genus level was generated using the Phyloseq package.

We chose least absolute shrinkage and selection operator (LASSO) regression as a modeling method to predict wheat quality for the following reasons: i) to avoid overfitting, which may be problematic with other regression methods (least square regression or general linear model), especially when there are many explanatory variables, (ii) to be able to select only the most important predictive variables (i.e., feature), to reduce the mean error of the model, and (iii) to have an interpretable model. The microbial features included: principal components 1-5 derived from the microbial OTU and carbon usage tables, the abundance of N-cycle related gene, the F:B ratio, and the bacterial and fungal alpha-diversity. First, we standardized the data (other than the PCs) using the *scale* function and then selected the optimal lambda values with 10-fold cross validation. We selected the best penalty score based on the lowest lambda value, which indicates non-collinear effects and low levels of inflated variance in the selected variables. Then, we evaluated the model accuracy and performance using the best lambda values. The predictive strength of the best LASSO model for grain quality was tested using the *prediction* function of the caret package. The Akaike Information Criterion (AIC) and the Bayesian Information Criterion (BIC) were also calculated to evaluate the models’ performance. Finally, we compared the accuracy and performance across the different sampling dates.

## Results

### Effect of experimental treatments on microbial parameters

The sampling date significantly affected all microbial parameters, including microbial carbon utilization, microbial diversity, the F:B ratio, and the abundance of N-cycle-related genes (Tables 1 and 2). Furthermore, the structure of the bacterial and archaeal community was influenced by sampling dates and blocks whereas the fungal community was influenced by sampling dates, block, and wheat genotypes (Table 2). The alpha diversity of microbial communities (16S and ITS) was not significantly affected by the precipitation exclusion treatments and wheat genotypes (P >0.05). However, significant differences in the Shannon (P<0.001) and Simpson (P < 0.001) diversity indices, as well as the phylogenetic diversity (P <0.001) of the bacterial and archaeal, and fungal communities were observed among sampling dates. There was a significant interactive effect (P < 0.05) of the precipitation treatment and wheat genotype on the abundance of archaeal *amoA, nirK* and *nosZ* genes, and a significant difference in the F: B ratio across sampling dates (P < 0.001) (Table 1).

**Table 1.** Three-way repeated measure ANOVA for bacterial and archaeal ammonia monooxygenase, nitrite reductase, nitrous oxide reductase gene abundance and the 16S: ITS genes ratio for the effect of precipitation exclusion treatments, sampling dates and genotype.

|  | *AOA* | *AOB* | *nirK* | *nosZ* | *F:B ratio* |
| --- | --- | --- | --- | --- | --- |
| treatment | 1.449 | 0.241 | 0.940 | 1.027 | 0.467 |
| date | 46.382*** | 40.379*** | 40.176*** | 79.707*** | 86.755*** |
| genotype | 0.205 | 0.006 | 0.388 | 0.689 | 0.043 |
| block | 2.180* | 3.175** | 2.682* | 0.995 | 0.918 |
| treatment × genotype | 4.782** | 0.993 | 4.356** | 3.188** | 0.854 |

F-values are shown in the table.
Treatment: treatments with precipitation exclusion (0%, 25%, 50%, 75%). Date: sampling dates. Genotype: drought-sensitive wheat and drought-tolerant wheat. ANOVA significance, “.” $0.1 < P < 0.05$ ; “\*” $P < 0.05$ ; “\*\*” $P < 0.01$ ; “\*\*\*” $P < 0.001$

**Table 2.** Permanova based on Bray Curtis dissimilarities for microbial carbon utilization profiling (Biolog EcoPlate) and community structure based on 16S rRNA gene and ITS region amplicon for the effect of precipitation exclusion treatments, sampling dates and genotype.

|  | **Biolog** |  |  | **16S** |  |  | **ITS** |  |  |
| --- | --- | --- | --- | --- | --- | --- | --- | --- | --- |
|  | R<sup>2</sup> | F | Pr(>F) | R<sup>2</sup> | F | Pr(>F) | R<sup>2</sup> | F | Pr(>F) |
| treatment | 0.013 | 4.95 | 0.002** | 0.003 | 0.95 | 0.419 | 0.004 | 1.45 | 0.086 |
| date | 0.105 | 39.71 | 0.001*** | 0.01 | 5.06 | 0.001*** | 0.01 | 2.90 | 0.001*** |
| genotype | 0.002 | 0.58 | 0.754 | 0.00 | 1.04 | 0.29 | 0.01 | 2.61 | 0.003** |
| block | 0.005 | 1.83 | 0.108 | 0.01 | 4.36 | 0.001*** | 0.03 | 11.72 | 0.001*** |
| genotype× treatment | 0.002 | 0.81 | 0.506 | 0.00 | 1.51 | 0.061 | 0.00 | 1.71 | 0.039* |

Treatment: precipitation exclusion (0%, 25%, 50%, 75%). Date: sampling dates. Genotypes: drought-sensitive wheat and drought-tolerant wheat. “.” 0.1 < P < 0.05; “\*” P < 0.05; “\*\*” P < 0.01; “\*\*\*” P < 0.001

### Correlation between microbial and grain quality parameters

We did not find a significant effect of rainfall exclusion treatment on grain qualities but found a significant effect of wheat genotype on protein content (P<0.001) and PMT (P<0.001), so we decided to treat the two genotypes separately and all the precipitation treatments together. Correlations between grain quality and microbial carbon use fluctuated over time (Table 3). The carbon sources were all negatively correlated to grain quality indicators for the DT genotype whereas both positive and negative correlations were found for the DS genotype (Table 3). The absolute abundance of microbial N-cycling genes was found to be correlated to grain quality measurements for soil collected on the early sampling dates (Table 4). The *amoA* (archaeal and bacterial), *nirK* and *nosZ* genes quantified in the DT genotype samples on May 10 and May 24 were negatively correlated to protein and gluten content (Table 4). Only the F:B ratio was positively correlated to protein content (Table 4). For the DS genotype, the *amoA* (archaeal and bacterial) and the *nosZ* genes were negatively correlated to the grain quality parameters and the F:B ratio was positively correlated to PMT for soil samples collected on May 24 (Table 4). Many significant correlations between microbial richness/diversity indices and grain baking quality were found, mostly for the DT genotype (Table 5). Significant correlations between microbial community descriptors (PCA axes for OTUs and microbial carbon use) and grain quality indicators for sampling dates in May and June were also identified.

**Table 3:**
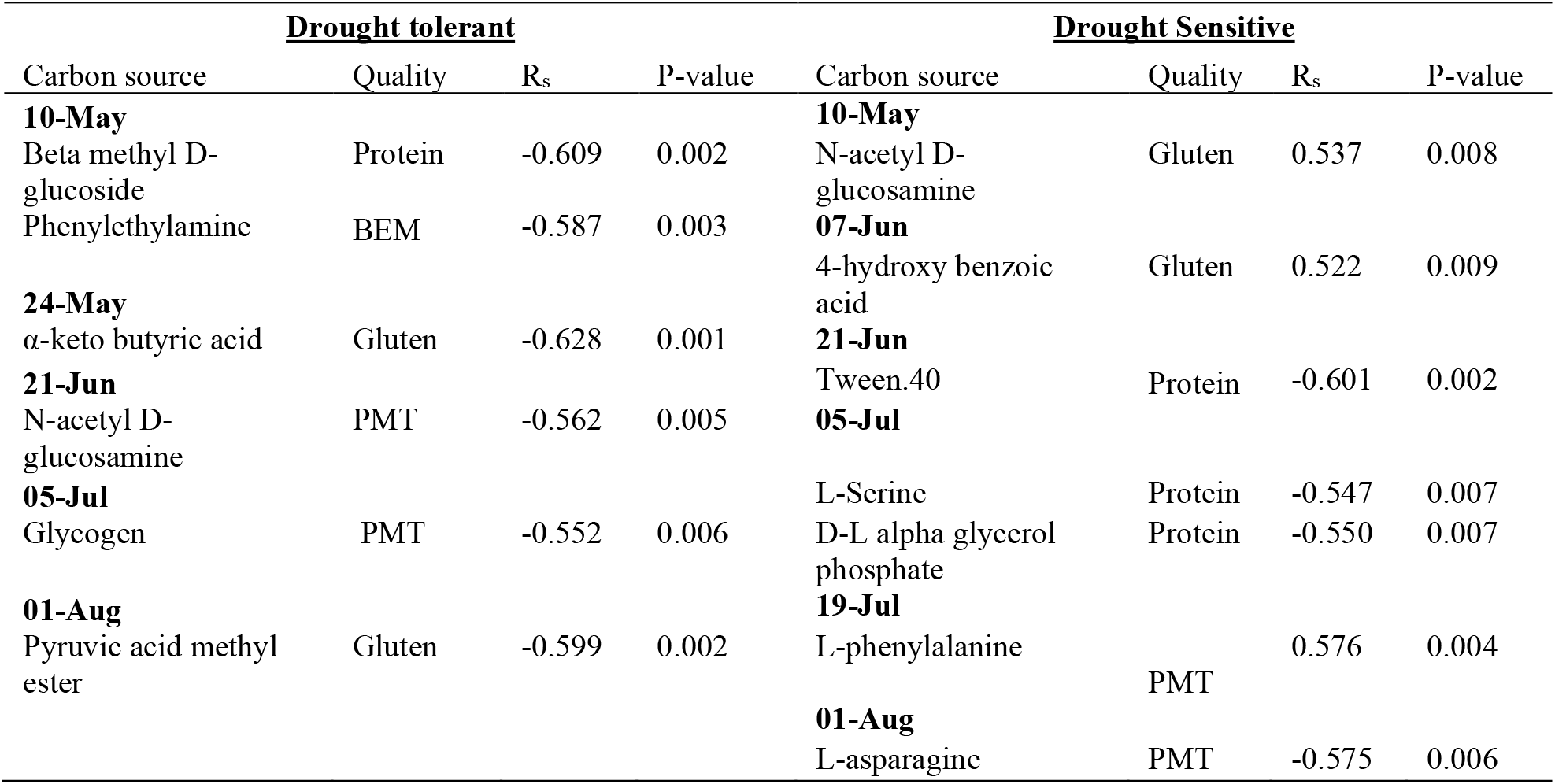
Significant (P<0.05) Spearman correlations between microbial carbon utilization and grain baking quality for each sampling date (N=24).

**Table 4:** Significant (P<0.05) Spearman correlations between functional gene abundance and grain baking quality for each sampling dates (N=24).

| Drought tolerant |  |  |  | Drought sensitive |  |  |  |
| --- | --- | --- | --- | --- | --- | --- | --- |
| Gene | Quality | R<sub>s</sub> | P-value | Gene | Quality | R<sub>s</sub> | P-value |
| 10-May |  |  |  | 24-May |  |  |  |
| *nosZ* | Gluten | -0.406 | 0.054 | *AOB* | Gluten | -0.504 | 0.012 |
| 24-May |  |  |  | *AOA* | Protein | -0.406 | 0.055 |
| *AOB* | Gluten | -0.450 | 0.031 | *nosZ* | BEM | -0.400 | 0.059 |
| *nirK* | Protein | -0.441 | 0.035 | *F:B ratio* | PMT | 0.425 | 0.043 |
| *AOA* | Protein | -0.578 | 0.004 | 07-Jun |  |  |  |
| *F: B Ratio* | Protein | 0.547 | 0.007 | *nirK* | Gluten | -0.441 | 0.035 |
| 07-Jun |  |  |  | 21-Jun |  |  |  |
| *F: B Ratio* | Protein | 0.426 | 0.048 | *F: B Ratio* | Protein | 0.406 | 0.054 |
| 21-Jun |  |  |  | *F: B Ratio* | PMT | -0.406 | 0.055 |
| *AOA* | Protein | -0.563 | 0.005 | *F: B Ratio* | BEM | 0.492 | 0.017 |
| *AOA* | PMT | 0.404 | 0.056 | 19-Jul |  |  |  |
| 05-Jul |  |  |  | *nirK* | Gluten | 0.558 | 0.009 |
| *nirK* | Gluten | -0.443 | 0.034 |  |  |  |  |
| *nosZ* | PMT | 0.401 | 0.058 |  |  |  |  |
| *F: B Ratio* | BEM | 0.479 | 0.021 |  |  |  |  |
| 19-Jul |  |  |  |  |  |  |  |
| *AOA* | Protein | -0.426 | 0.042 |  |  |  |  |
| 01-Aug |  |  |  |  |  |  |  |
| *nosZ* | PMT | 0.392 | 0.058 |  |  |  |  |

**Table 5:**
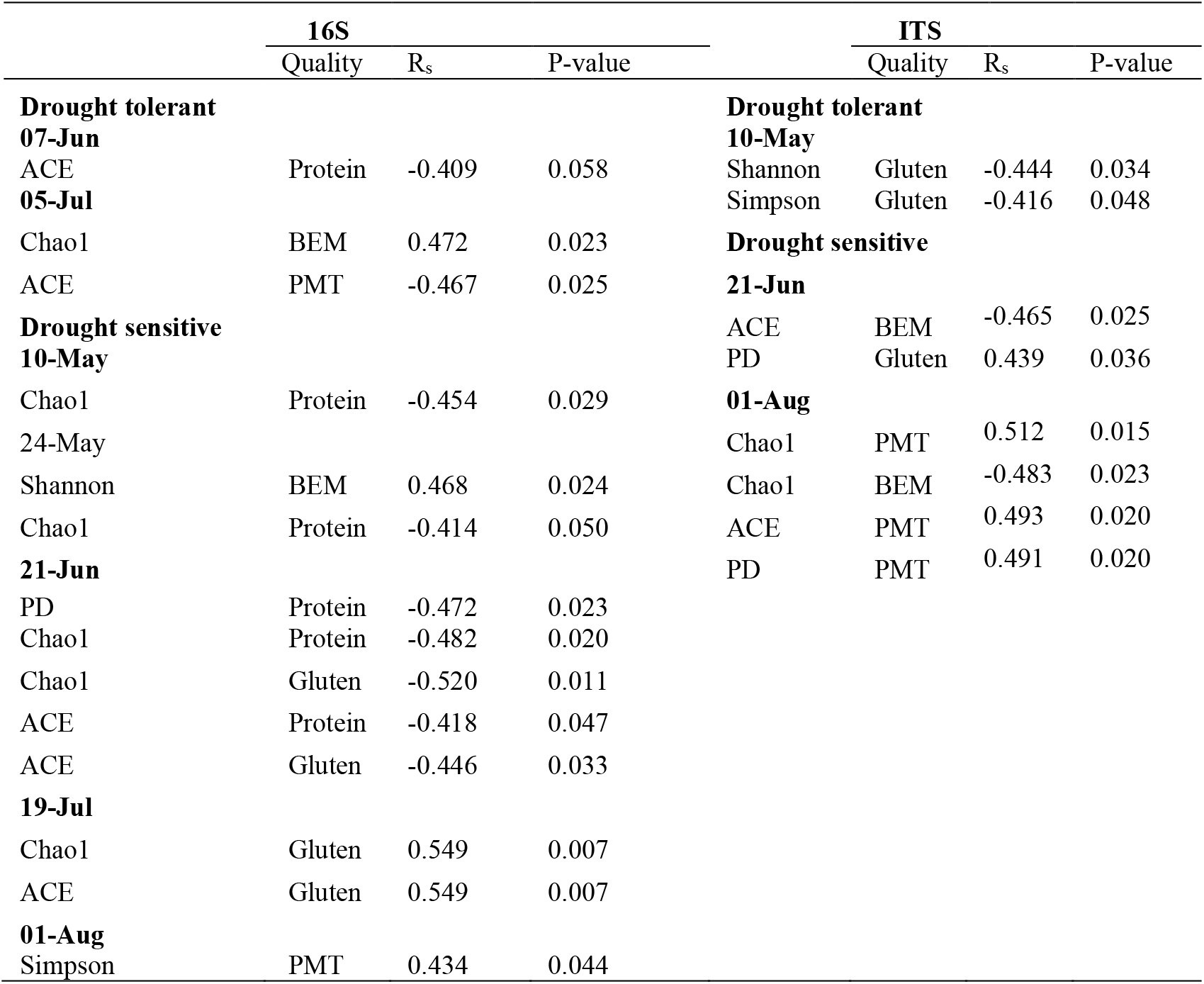
Significant (P<0.05) Spearman correlations between bacterial and archaeal and fungal richness and diversity and grain baking quality for each sampling dates (N=24).

|  | 16S |  |  |  | ITS |  |  |
| --- | --- | --- | --- | --- | --- | --- | --- |
|  | Quality | R<sub>s</sub> | P-value |  | Quality | R<sub>s</sub> | P-value |
| **Drought tolerant** |  |  |  | **Drought tolerant** |  |  |  |
| **07-Jun** |  |  |  | **10-May** |  |  |  |
| ACE | Protein | -0.409 | 0.058 | Shannon | Gluten | -0.444 | 0.034 |
| **05-Jul** |  |  |  | Simpson | Gluten | -0.416 | 0.048 |
| Chao1 | BEM | 0.472 | 0.023 | **Drought sensitive** |  |  |  |
| ACE | PMT | -0.467 | 0.025 | **21-Jun** |  |  |  |
| **Drought sensitive** |  |  |  | ACE | BEM | -0.465 | 0.025 |
| **10-May** |  |  |  | PD | Gluten | 0.439 | 0.036 |
| Chao1 | Protein | -0.454 | 0.029 | **01-Aug** |  |  |  |
| 24-May |  |  |  | Chao1 | PMT | 0.512 | 0.015 |
| Shannon | BEM | 0.468 | 0.024 | Chao1 | BEM | -0.483 | 0.023 |
| Chao1 | Protein | -0.414 | 0.050 | ACE | PMT | 0.493 | 0.020 |
| **21-Jun** |  |  |  | PD | PMT | 0.491 | 0.020 |
| PD | Protein | -0.472 | 0.023 |  |  |  |  |
| Chao1 | Protein | -0.482 | 0.020 |  |  |  |  |
| Chao1 | Gluten | -0.520 | 0.011 |  |  |  |  |
| ACE | Protein | -0.418 | 0.047 |  |  |  |  |
| ACE | Gluten | -0.446 | 0.033 |  |  |  |  |
| **19-Jul** |  |  |  |  |  |  |  |
| Chao1 | Gluten | 0.549 | 0.007 |  |  |  |  |
| ACE | Gluten | 0.549 | 0.007 |  |  |  |  |
| **01-Aug** |  |  |  |  |  |  |  |
| Simpson | PMT | 0.434 | 0.044 |  |  |  |  |

### Model performance in predicting grain quality at different plant growth stages

We applied least absolute shrinkage and selection operator (LASSO) regressions for each sampling date separately, to identify the date where model accuracy would be maximal to predict grain quality. In the case of the DT genotype, we obtained a high model accuracy to predict certain grain quality indicators, with mean square errors ranging from 0.08 to 0.51, and AIC values inferior to −8.35 (Table 6). The best models identified were based on microbial indicators from May 10, May 24, and June 07. Gluten and protein content predicted with the LASSO regression had the highest accuracy for microbial indicators measured from samples collected on May 10. These models selected 11 and 8 variables, resulting in R^2^ of 0.95 and 0.76, for gluten and protein respectively (Table 6 and Figure 1). These models were cross validated with lambda λ values of 0.04 to 0.15, which resulted in the lowest mean square errors (0.08 to 0.36). The model accuracy for gluten and protein prediction decreased over time, and it was even impossible to generate a significant model for some dates (Table 6). The best sampling dates for the other quality indicators were later in the growing season with June 7 being the best sampling time to predict BEM (R^2^=0.92) and May 24 being the best time to predict PMT (R^2^=0.57) (Table 6 and Fig. 1). The most parsimonious model across all indicators was the one predicting PMT which only included 2 predictors (Table 6). For some sampling dates, no descriptive variable in the microbial dataset was selected by the LASSO procedure, resulting in a null model (Table 6). This was the case for gluten on June 7, July 19, and August 1, for PMT on May 10, June 7, June 21, July 5 and July 19, and for BEM on June 21 and July 05 (Table 6).

**Table 6:** Comparative analysis of the LASSO model performance for the wheat grain quality of the drought-tolerant genotype (DT).

|  | T1 | T2 | T3 | T4 | T5 | T6 | T7 |
| --- | --- | --- | --- | --- | --- | --- | --- |
| Date | 10-May | 24-May | 07-Jun | 21-Jun | 05-Jul | 19-Jul | 01-Aug |
| Gluten (DT) |  |  |  |  |  |  |  |
| C.V (best Lambda) | 0.04 | 0.38 |  | 0.56 | 0.72 |  |  |
| AIC | -16.14 | 2.00 |  | 1.33 | 2.00 |  |  |
| BIC | -15.09 | 3.04 |  | 2.42 | 3.09 |  |  |
| Nb of variables: | 11 |  |  | 1 | 1 |  |  |
| MSE (Mean Square Error) | 0.08 | 0.95 |  | 0.92 | 0.95 |  |  |
| R<sup>2</sup> | **0.95** | 0.15 |  | 0.54 | 0.54 |  |  |
| Protein (DT) |  |  |  |  |  |  |  |
| C.V (best Lambda) | 0.15 | 0.24 | 0.19 | 0.28 | 0.46 | 0.36 | 0.18 |
| AIC | -11.64 | -9.21 | -9.56 | -3.40 | 2.00 | 2.00 | -8.24 |
| BIC | -10.51 | -8.07 | -8.47 | -2.26 | 3.14 | 3.14 | -7.15 |
| Nb of variables: | 8 | 5 | 7 | 2 | 1 | 1 | 2 |
| MSE (Mean Square Error) | 0.36 | 0.47 | 0.43 | 0.73 | 0.96 | 0.96 | 0.53 |
| R<sup>2</sup> | **0.76** | 0.69 | 0.72 | 0.33 | 0.22 | 0.14 | 0.57 |
| PMT (DT) |  |  |  |  |  |  |  |
| C.V (best Lambda) |  | 0.21 |  |  |  |  | 0.42 |
| AIC |  | -8.35 |  |  |  |  | 2.00 |
| BIC |  | -7.21 |  |  |  |  | 3.18 |
| Nb of variables: |  | 2 |  |  |  |  | 1 |
| MSE (Mean Square Error) |  | 0.51 |  |  |  |  | 0.96 |
| R<sup>2</sup> |  | **0.57** |  |  |  |  | 0.19 |
| BEM (DT) |  |  |  |  |  |  |  |
| C.V (best Lambda) | 0.38 | 0.25 | 0.03 |  |  | 0.14 | 0.20 |
| AIC | -2.34 | -7.31 | -17.00 |  |  | -8.92 | -5.32 |
| BIC | -1.21 | -6.17 | -15.91 |  |  | -7.78 | -4.14 |
| Nb of variables: | 1 | 2 | 10 |  |  | 7 | 4 |
| MSE (Mean Square Error) | 0.77 | 0.55 | 0.09 |  |  | 0.48 | 0.65 |
| R<sup>2</sup> | 0.35 | 0.50 | **0.92** |  |  | 0.58 | 0.47 |

AIC= Akaike Information Criterion
BIC=Bayesian Information Criterion
C. V= Cross validation

**Figure 1.**
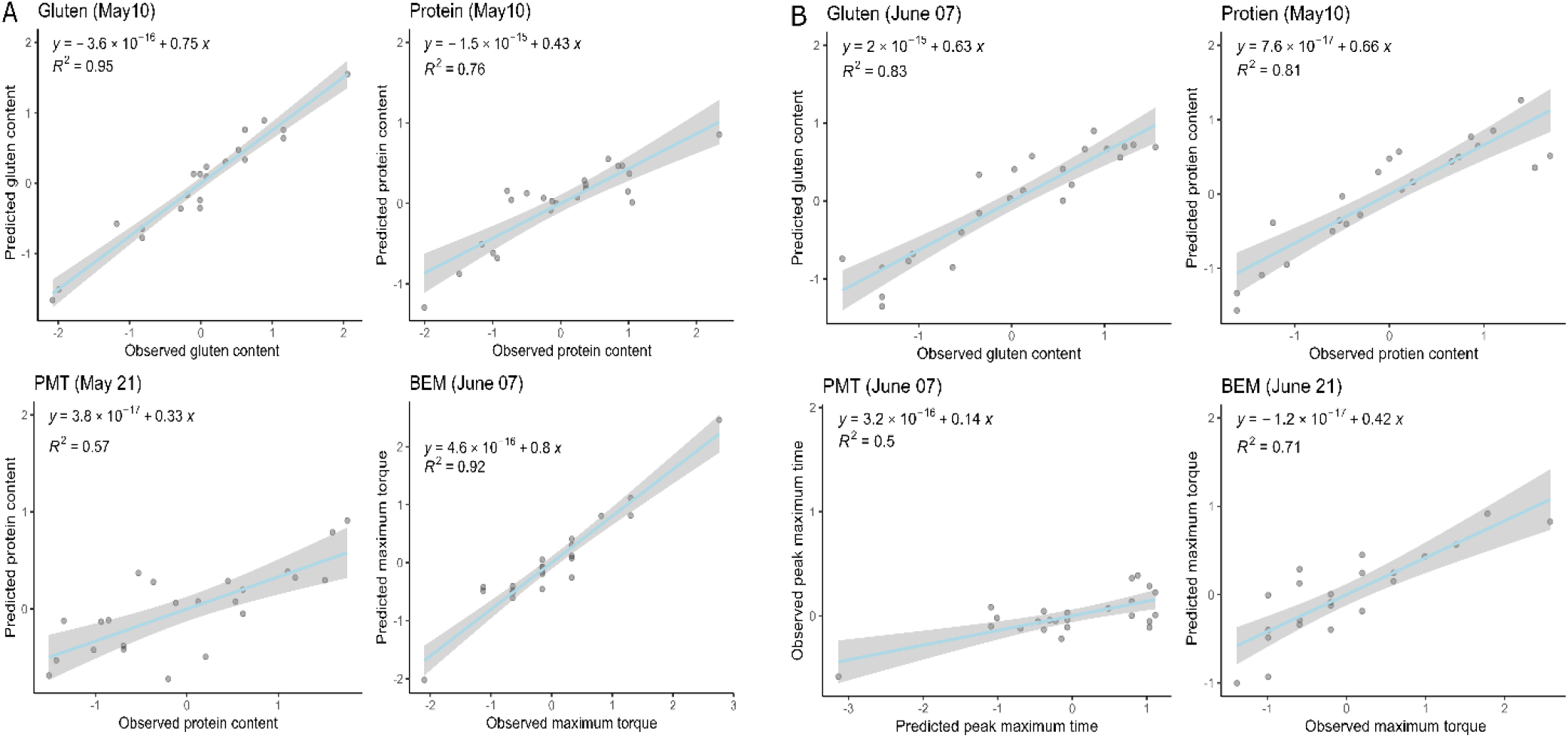
Observed values vs. predicted values from LASSO regression models for wheat grain gluten and protein content and flour maximum torque (BEM) and peak maximum time (PMT) for the drought-tolerant (A) and drought-sensitive genotypes (B).

The overall model performance in predicting grain quality for the DS genotype was lower than the DT genotype (Table 7). Maximum accuracy of LASSO regression model was observed on June 7 for gluten and PMT, on May 10 for protein, and June 21 for BEM (Table 7). The best PMT and BEM predictive models used about half the number of the total predictors used in the best gluten and protein predictive models (PMT: 4, BEM: 6, gluten: 14 and protein: 11) (Table 7 and 9). These models included many fungal indicators (Table 9). Predictive modeling of protein content between May 24 and July 05, and on August 1 was unsuccessful and the level of accuracy of the model weas lower on July 19 (Table 7). A similar trend was observed for PMT: sampling dates after June 7 resulted in less accurate or no model at all (Table 7). BEM prediction was also unsuccessful for samples collected on June 07. Similar to the DT genotype, the predictive models for the DS genotype dataset showed the best accuracy for quality prediction with microbial data from the May and June samplings.

**Table 7:** Comparative analysis of the model performance of LASSO for the wheat grain quality of drought-sensitive genotype (DS).

|  | T1 | T2 | T3 | T4 | T5 | T6 | T7 |
| --- | --- | --- | --- | --- | --- | --- | --- |
| Date | 10-May | 24-May | 07-Jun | 21-Jun | 05-Jul | 19-Jul | 01-Aug |
| **Gluten (DS)** |  |  |  |  |  |  |  |
| AIC | -8.28 | 2.00 | -16.01 | -14.60 | -1.97 | -14.66 | 0.67 |
| BIC | -7.14 | 3.14 | -14.84 | -13.46 | -0.83 | -13.52 | 1.76 |
| C.V (best Lambda) | 0.18 | 0.45 | 0.08 | 0.09 | 0.34 | 0.10 | 0.32 |
| Nb of variables: | 6 | 1 | 14 | 11 | 1 | 10 | 1 |
| MSE (Mean Square Error) | 0.51 | 0.96 | 0.21 | 0.23 | 0.78 | 0.23 | 0.89 |
| R<sup>2</sup> | 0.61 | 0.22 | 0.83 | 0.81 | 0.30 | 0.81 | 0.17 |
| **Protein (DS)** |  |  |  |  |  |  |  |
| C.V (best Lambda) | 0.06 |  |  |  |  | 0.17 |  |
| AIC | -15.15 |  |  |  |  | -7.69 |  |
| BIC | -14.02 |  |  |  |  | -6.55 |  |
| Nb of variables: | 11 |  |  |  |  | 6 |  |
| MSE (Mean Square Error) | 0.21 |  |  |  |  | 0.54 |  |
| R<sup>2</sup> | 0.81 |  |  |  |  | 0.53 |  |
| **PMT (DS)** |  |  |  |  |  |  |  |
| C.V (best Lambda) | 0.19 | 0.41 | 0.33 |  | 0.38 |  | 0.24 |
| AIC | -5.76 | -3.94 | -3.56 |  | 2.00 |  | -1.55 |
| BIC | -4.63 | -2.80 | -2.38 |  | 3.14 |  | -0.46 |
| Nb of variables: | 4 | 1 | 4 |  | 1 |  | 1 |
| MSE (Mean Square Error) | 0.62 | 0.70 | 0.73 |  | 0.96 |  | 0.79 |
| R<sup>2</sup> | 0.35 | 0.45 | 0.50 |  | 0.15 |  | 0.24 |
| **BEM (DS)** |  |  |  |  |  |  |  |
| C.V (best Lambda) | 0.13 | 0.36 |  | 0.19 | 0.18 | 0.13 | 0.17 |
| AIC | -10.11 | -0.70 |  | -10.37 | -8.21 | -8.67 | 2.00 |
| BIC | -9.02 | 0.39 |  | -9.28 | -7.11 | -7.58 | 3.04 |
| Nb of variables: | 11 | 2 |  | 6 | 5 | 4 | 1 |
| MSE (Mean Square Error) | 0.40 | 0.83 |  | 0.39 | 0.49 | 0.47 | 0.95 |
| R<sup>2</sup> | 0.65 | 0.32 |  | 0.71 | 0.61 | 0.56 | 0.03 |

AIC= Akaike Information Criterion
BIC=Bayesian Information Criterion
C. V= Cross validation

### Microbial features selected in the optimal models

The best LASSO models for the DT genotype contained microbial features that varied but were often the principal components derived from OTU tables or carbon utilization patterns, or the alpha diversity indices. Bacterial and archaeal OTUs from the *Nitrosphaera, Rhodoplanes, Solirubrobacter*, and *Terrimicrobium* were the main contributors to the principal component 2 (explained variance: 5.1%) calculated from the May 10 dataset that was selected in the models for gluten and protein content (Fig. 2). In contrast, the main contributors to the bacterial and archaeal principal component 1 (explained variance: 6.0%), 2 (5.2%) and 3 (5.1%) selected for the model predicting BEM on June 7 were from the *Conexibacter, Gaiella, Nitrososphaera, Hyphomicrobium* and *Gp16* genera (Fig. 2). The fungal OTUs that contributed to the principal components selected in the May and June models belonged to the *Mortierella, Ganoderma*, and *Giliomastix* genera (Fig. 2). We found a negative relationship between the fungal phylogenetic diversity index and gluten content and a positive relationship between bacterial Simpson diversity and gluten content and BEM in the May 10 and June 7 models (Table 8).

**Figure 2.**
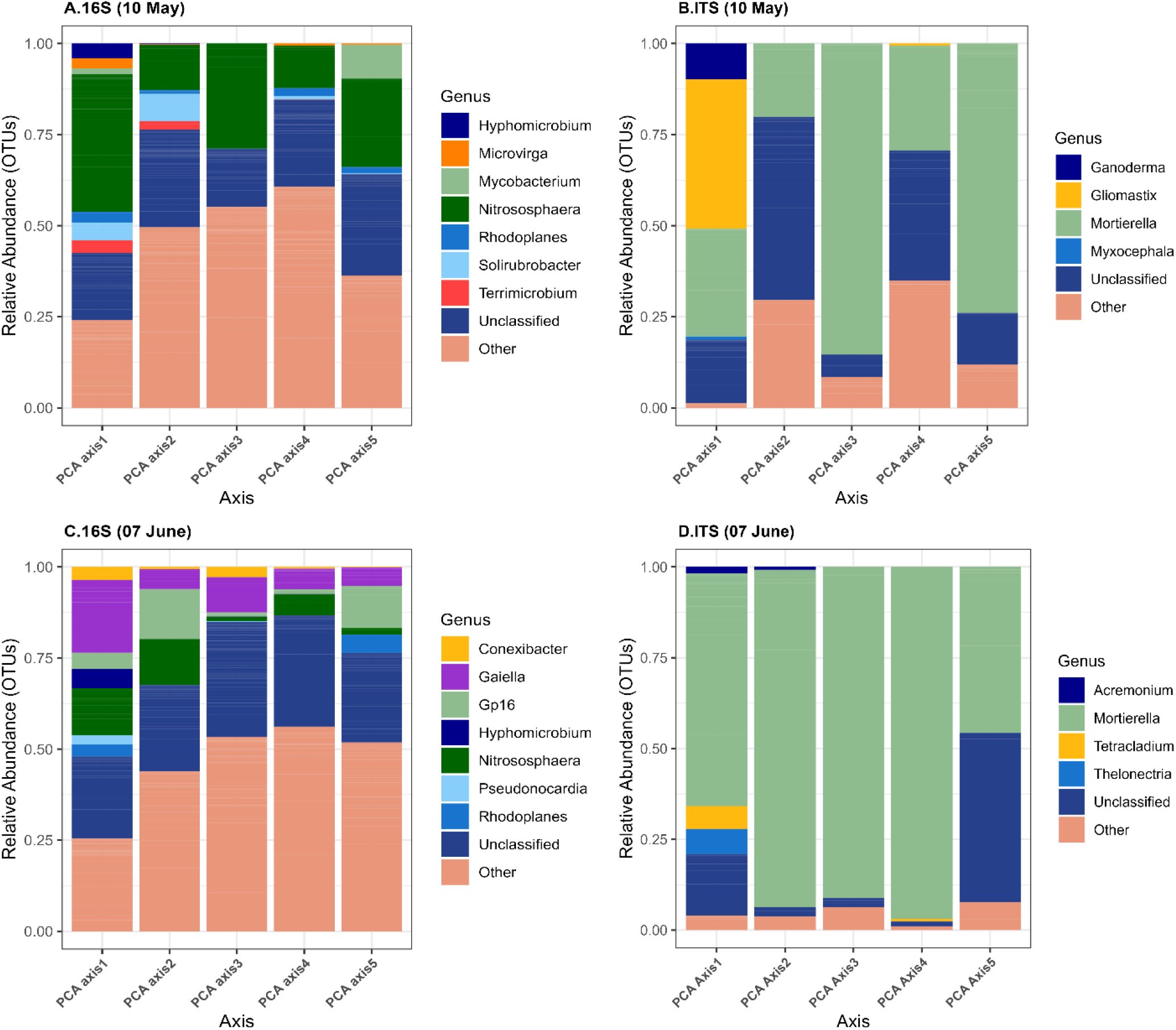
The relative abundance of the bacterial and archaeal (A, C) and fungal (B, D) genera significantly correlated with the first five principal components for the drought tolerant genotype for the May 10 (A, B) and June 7 (C, D) sampling dates. Others: various genera with relative abundances below 0.1%.

**Table 8:**
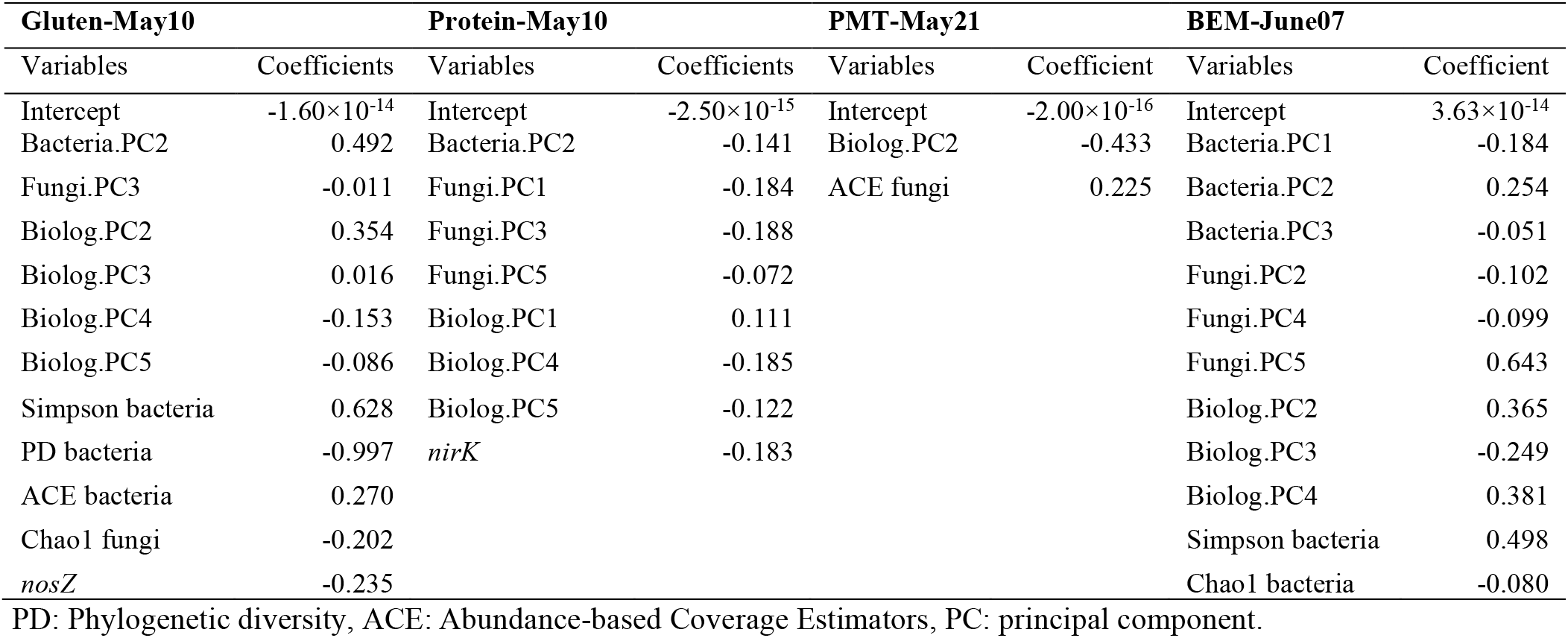
Microbial parameters included in the LASSO models for wheat grain quality of the drought-tolerant genotype (DT).

| Gluten-May10 |  | Protein-May10 |  | PMT-May21 |  | BEM-June07 |  |
| --- | --- | --- | --- | --- | --- | --- | --- |
| Variables | Coefficients | Variables | Coefficients | Variables | Coefficient | Variables | Coefficient |
| Intercept | $-1.60 \times 10^{-14}$ | Intercept | $-2.50 \times 10^{-15}$ | Intercept | $-2.00 \times 10^{-16}$ | Intercept | $3.63 \times 10^{-14}$ |
| Bacteria.PC2 | 0.492 | Bacteria.PC2 | -0.141 | Biolog.PC2 | -0.433 | Bacteria.PC1 | -0.184 |
| Fungi.PC3 | -0.011 | Fungi.PC1 | -0.184 | ACE fungi | 0.225 | Bacteria.PC2 | 0.254 |
| Biolog.PC2 | 0.354 | Fungi.PC3 | -0.188 |  |  | Bacteria.PC3 | -0.051 |
| Biolog.PC3 | 0.016 | Fungi.PC5 | -0.072 |  |  | Fungi.PC2 | -0.102 |
| Biolog.PC4 | -0.153 | Biolog.PC1 | 0.111 |  |  | Fungi.PC4 | -0.099 |
| Biolog.PC5 | -0.086 | Biolog.PC4 | -0.185 |  |  | Fungi.PC5 | 0.643 |
| Simpson bacteria | 0.628 | Biolog.PC5 | -0.122 |  |  | Biolog.PC2 | 0.365 |
| PD bacteria | -0.997 | *nirK* | -0.183 |  |  | Biolog.PC3 | -0.249 |
| ACE bacteria | 0.270 |  |  |  |  | Biolog.PC4 | 0.381 |
| Chao1 fungi | -0.202 |  |  |  |  | Simpson bacteria | 0.498 |
| *nosZ* | -0.235 |  |  |  |  | Chao1 bacteria | -0.080 |

PD: Phylogenetic diversity, ACE: Abundance-based Coverage Estimators, PC: principal component.

Principal components derived from carbon utilization patterns were also included in all our most accurate models for the DT genotype (Table 8). The models predicting protein and gluten content (May 10) selected 4 of the top 5 principal components included, for which the most important contributing carbon substrates were Putrescine (r_s_=-0.91; P<0.001), L-Arginine (r_s_=0.74; P <0.001), Pyruvic Acid methyl ester (r_s_=-0.62; P<0.001), Glycogen (r_s_=0.59; P<0.001) and L-Threonine (r_s_= −0.56; P<0.001). The model predicting BEM (June 7) selected principal component 2 (explained variance: 9.3%), 3 (7.4%), and 4 (4.7%) and the most important contributing carbon substrates of the principle components were alpha-cyclodextrin (r_s_=0.69; P=0.002), alpha-keto butyric Acid (r_s_=0.68; P=0.003), γ-amino butyric acid (r_s_= −0.66; P =0.006), Glucose 1-phosphate (r_s_=-0.71; P=0.001). Finally, the principal component 2 (explained variance: 7.5%) selected in the model predicting PMT (May 24) was correlated to glycogen (r_s_=0.59; P=0.002), alpha-cyclodextrin (r_s_=0.68; P<0.001) and γ-amino butyric acid (r_s_=-0.65; P=0.004). We also observed a negative relationship between protein content and *nirK* (regression coef. = −0.183) and gluten content and *nosZ* (regression coef. = −0.235) in the models obtained on May 10 (Table 7).

As for the DT genotype models, the models for the DS genotype were mainly composed of principal components calculated from the OTU tables and from the carbon utilization patterns, and from alpha-diversity indices (Table 9). The LASSO model predicting protein content selected the bacterial principal component 4 (explained variance: 4.9%) for the May 10 sampling date (Table 9). This principal component was correlated with OTUs belonging to the *Nitrososphaera, Rhodoplanes, Solirubrobacter*, and *Terricomicrobium* (Fig. 3). On the same date, the fungal OTUs contributing the most to the principal component 1 (explained variance: 7.3%), 3 (5.6%), 4 (5.5%), and 5 (5.2%) belonged to the *Acremonium, Mortierella, Pezizella*, and *Tetracladium* (Fig. 3). On June 7, the models predicting gluten content and PMT selected the bacterial principal components 2, 4, and 5 (Table 9). These axes explained 4.7-4.5% of the variation and were correlated to OTUs related to *Giella, Gp6, Hyphomicrobium, Nitrososphaera, Rhodoplanes*, and *Solirubrobacter* (Fig. 3). On June 21, the model predicting BEM selected the bacterial principal components 2 and 4 (Table 9), which explained 4.8% and 4.7% of the variation and were correlated to OTUs related to *Nitrososphaera, Giella, Gp6, Pseudonocardia, Bradyrhizobium*, and *Lysinibacillus* (Table 9). The fungal PC 1 (7%), 2 (6.2%), 4 (5.0%), and 5 (4.9%) selected for the June 7 were correlated to OTUs related to *Acremonium, Mortierella*, and *Tetracladium* (Fig. 3). The fungal PC2 (5.8%) selected in the model for BEM in June 21 was linked to OTUs related to *Ganoderma, Mortierella, Pezizella* and *Pseudeurotium*. The protein and BEM models of the DS genotype selected the fungal Chao1 index, which positively influenced the quality whereas the gluten and protein models selected fungal phylogenetic diversity indices that positively affected gluten content on June 07, and negatively affected protein content on May 10 (Table 9).

**Table 9:** Microbial parameters included in the LASSO models for the wheat grain quality of the drought-sensitive genotype (DS).

| Gluten-June 07 |  | Protein-May10 |  | PMT-June 07 |  | BEM-June21 |  |
| --- | --- | --- | --- | --- | --- | --- | --- |
| Variables | Coefficients | Variables | Coefficients | Variables | Coefficients | Variables | Coefficients |
| Intercept | $7.96 \times 10^{-15}$ | Intercept | $-1.80 \times 10^{-17}$ | Intercept | $3.57 \times 10^{-16}$ | Intercept | $-8 \times 10^{-17}$ |
| Bacteria.PC2 | -0.018 | Bacteria.PC4 | -0.541 | Bacteria.PC4 | -0.009 | Bacteria.PC2 | -0.010 |
| Bacteria.PC5 | 0.216 | Fungi.PC1 | -0.219 | Fungi.PC4 | 0.146 | Bacteria.PC4 | -0.263 |
| Fungi.PC1 | 0.012 | Fungi.PC3 | -0.093 | Fungi.PC5 | 0.120 | Fungi.PC2 | -0.086 |
| Fungi.PC2 | -0.361 | Fungi.PC4 | -0.262 | Biolog.PC5 | 0.026 | Biolog.PC4 | -0.151 |
| Fungi.PC4 | -0.026 | Fungi.PC5 | 0.141 |  |  | Chao1 fungi | 0.182 |
| Fungi.PC5 | -0.072 | Biolog.PC3 | 0.027 |  |  | F:B ratio | 0.213 |
| Biolog.PC1 | 0.317 | Biolog.PC4 | -0.099 |  |  |  |  |
| Biolog.PC3 | -0.078 | Biolog.PC5 | -0.009 |  |  |  |  |
| Biolog.PC5 | 0.024 | Chao1 bacteria | -0.284 |  |  |  |  |
| Simpson bacteria | 0.089 | Chao1 fungi | 0.154 |  |  |  |  |
| PD fungi | -0.150 | PD fungi | 0.012 |  |  |  |  |
| AOB | -0.460 |  |  |  |  |  |  |
| *nosZ* | 0.115 |  |  |  |  |  |  |
| F:B ratio | -0.031 |  |  |  |  |  |  |

PD: Phylogenetic diversity, F:B: Fungal: Bacterial ratio, PC: principal component.

**Figure 3.**
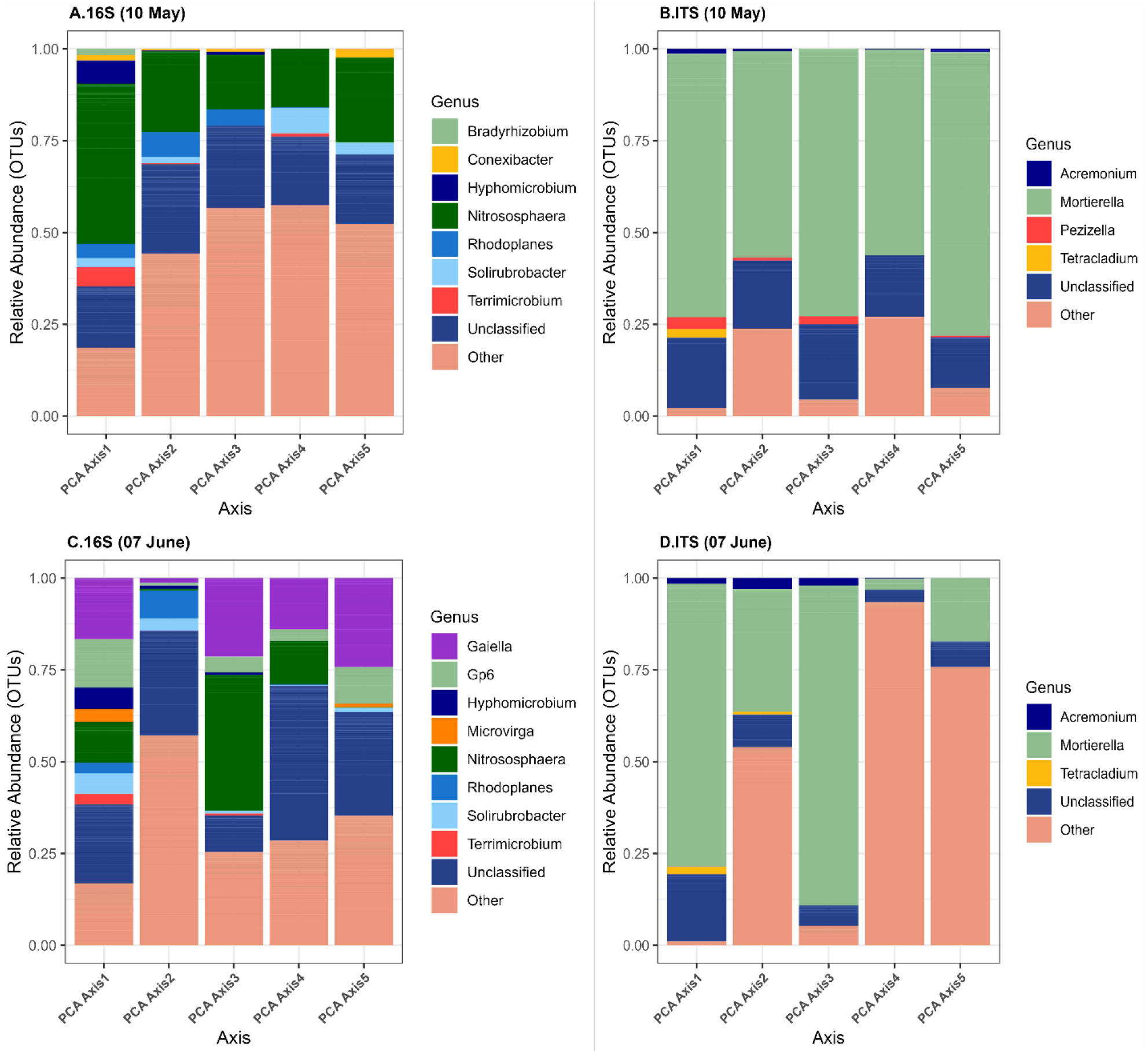
The relative abundance of the bacterial and archaeal (A, C) and fungal (B, D) genera significantly correlated with the first five principal components for the drought sensitive genotype for the May 10 (A, B) and June 7 (C, D) sampling dates. Others: various genera with relative abundances below 0.1%.

For the May 10 model (protein), the carbon substrates contributing the most to the selected principal components were beta-methyl D-glucoside (r_s_=0.61; P=0.001), D-glucosamine acid (r_s_= −0.58; P=0.003), D-galactonic acid y-lactone (r_s_ =-0.53; P=0.008). For the June 7 models (gluten and PMT), the carbon substrates contributing the most to the selected PC were Glucose 1-phosphate (r_s_=0.81; P<0.001), D-galactonic acid y-lactone (r_s_=0.64; P=0.0005), 4-hydroxy benzoic acid (r_s_= −0.66; P=0.0005), 2-hydroxy benzoic acid (r_s_=0.56, P=0.003). Finally, for the June 21 model (BEM), the carbon substrates contributing the most to the selected PC were L-phenylalanine (r_s_= 0.55; P= 0.003) and alpha-cyclodextrin (r_s_=-0.49; P=0.011). We also observed that the models selected the fungal: bacterial ratio, which negatively influenced the gluten content on June 7 and positively influenced BEM on June 21. There was a negative relationship between the abundance of the bacterial *amoA* gene and gluten content, and a positive relationship between *nosZ* and gluten content on June 7 (Table 9).

## Discussion

Plant- and soil-associated microbial communities vary throughout the seasons/plant growth stages (Chaparro *et al*. 2013, 2014; Moroenyane *et al*. 2021; Azarbad *et al*. 2022; Azarbad *et al*. 2021; Wang *et al*. 2022) and it was unsure what was the best timing to create models to predict wheat grain quality. By sampling the same field every 2 weeks and measuring a wide range of microbial parameters, we were able to show with LASSO regression that the predictive value of microbial parameters is optimal during the earlier stages of wheat growth, at the seedling (May) or tillering stages (June). Many microbial parameters were consistently singled out by the regression models, which could allude to a mechanistic link between grain quality and the parameter identified, or simply to covariation between the microbial parameter and grain quality due to a third unmeasured parameter. Our work focused on wheat, and although it would be interesting to see if similar patterns apply to other crops, it is the first and necessary step to start building microbial-based predictive models for crop yields and quality.

All the best models were made with data collected before the end of June, which is at the early stages of wheat growth in Quebec. This is coherent with our previous results that showed that good predictive models could be made with soil samples taken in May or June (Yergeau et al. 2020; Asad et al. 2021) even though different sampling point were not compared. Other work done on willows showed that early microbial community composition could predict the potential of the trees to decontaminate soil or to survive (Bell *et al*. 2014; Yergeau *et al*. 2015). Navarro-Noya *et al*. (2022) showed that the complexity of microbial structure and diversity increases with maize development, and that the effect of agricultural practices on the soil microbiome was more evident at the early stages, which could explain why early microbial indicators performed better. This is encouraging for future work, as the ultimate goal of this type of predictive modeling is to have a tool that could be used to guide management strategies for farmers. Maximum usefulness will happen if indicators of yields or quality can be measured early, when it is still possible to intervene. It could be that the sampling dates highlighted are the ones that are the most critical for wheat grain quality, but for wheat, it is generally thought that the grain filling stage (around mid July in Quebec) is the most critical stage in term of N nutrition for high quality grain (Zörb *et al*. 2018). However, unless there is an unlikely massive microbial immigration, the microorganisms that can modulate or are indicative of soil N availability are already present in the soil early at seeding, and it is likely that their abundance and diversity at this stage could predict wheat grain quality. In fact, it was recently suggested that, because of their potential to be influenced by legacy and current environmental conditions, microbial communities act as multivariate integrators of the current and past physico-chemical conditions of their immediate environment, making them highly suitable predictors for ecosystem processes (Correa-Garcia *et al*. 2022).

Microbiome data have characteristics (sparsity, high dimensionality, zero-inflated) that often make them challenging to use in models. Here, we transformed the OTU and carbon utilization patterns tables using eigenvalue decomposition, namely principal component analysis, which reduces the dimension of the datasets to n-1 principal components that are orthogonal (not collinear) and ordered in decreasing order of variance explanation, moving from several thousands of descriptors to 23, in the case of the OTU tables. We further reduced the dimensionality by only utilizing the first 5 principal components in our LASSO regression, with the idea that these components contained a large part of the variation in the original dataset. One downside of this approach is that it makes the models less directly interpretable, with principal components being composite variable for many OTUs or carbon sources. However, using correlation analyses of individual OTUs with the principal components we were able to identify taxonomic groups and carbon sources that were linked with the principal components. We also used LASSO regression that selects of the most significant variables and shrinks the regression coefficient of the other variable to zero, generally producing parsimonious, highly interpretable models containing a few variables. Although non-parametric methods (neural network, random forest, support vector machine, etc.) could produce more accurate models, they are often less interpretable, meaning that the predictors influencing the output cannot be easily identifiable. Still, our models had high accuracy of 50-95%. The predictive performance of LASSO regression to predict biological characteristics from microbiome data was shown to be excellent for zero-inflated data such as microbial OTU count tables (Xiao *et al*. 2018; Dong *et al*. 2020). We also had good results using linear regression coupled with forward/backward selection with a preselection of individual OTUs that showed the strongest correlations with the predictors (Yergeau *et al*. 2020; Asad *et al*. 2021).

General community descriptors were often selected as the best explanatory variables in the models. Alpha diversity indices and eigenvectors (such as principal components) derived from microbial community structures are integrators of many parameters. Interestingly, it suggests that shallow sequencing to recover alpha and beta diversity patterns together with community level carbon utilization profiling would be sufficient to model wheat grain quality. Additionally, some specific microbial parameters were consistently singled out by the analyses. For example, the negative relationships between wheat quality and the abundance of the *nirK, nosZ* and bacterial *amoA* genes were well aligned with previous work (Yergeau *et al*. 2020; Asad *et al*.2021). The relative abundance of OTUs belonging to the ammonia-oxidizing archaea taxon *Nitrososphaera* were also highly correlated with many of the principal components selected in the models, and the abundance of both the archaeal and the bacterial *amoA* genes was often negatively correlated to quality parameters. This further suggests the key role of nitrification and denitrification in wheat grain quality, as proposed before (Yergeau *et al*. 2020; Asad *et al*. 2021; Wang *et al*.2022). Since grain quality is linked to its protein content, it is energetically more efficient for the plant to uptake ammonia, which can directly be incorporated into amino acids, whereas nitrate will need to be transformed back to ammonia (Beeckman et al., 2018). Nitrate uptake also requires more energy than ammonia uptake (Beeckman et al., 2018). Finally, nitrate is prone to leach and is a substrate for denitrification, which will lead to loss of nitrogen to the atmosphere. Manipulating or inhibiting the activity of these microbial guilds using, for instance, natural or artificial nitrification inhibitors may increase wheat grain quality. However, this strategy will need to be further studied to understand potential unwanted effects, as a common nitrification inhibitor, nitrapyrin, was shown to have off-target effects on the soil microbial community (Schmidt *et al*. 2022) and that nitrate stimulates lateral root elongation and affects various signaling pathways in the plant (Beeckman et al. 2018). Microbiome manipulation is still in its infancy and, because of ecological processes underlying community assembly, it will be a challenge (Agoussar & Yergeau, 2021). It is also unclear if microorganisms involved in nitrification and denitrification are sufficient indicators for accurate modeling of the grain quality, and, consequently, if solely targeting this group will result in the expected increase in grain quality. As our model showed, general community structure and diversity seem to also have a prime importance in determining wheat grain quality.

Our previous work showed that significant predictive models could be parametrized using microbial data measured early in the growing season, across a transect of more than 500 km (Asad et al. 2021). Here, we sought to confirm that early microbial measurements were optimal for such predictive models by focussing on a single field and sampling it every two weeks for a complete growing season. Taken together, the two studies confirm that our microbial-based models are effective at a large spatial scale and that they are optimally build using samples taken early in the season. Although we used a different modeling approach than previously, the selection of ammonia-oxidizers by the models was shared with our previous studies (Yergeau *et al*. 2020; Asad *et al*. 2021), suggesting a potential key role of this functional guild for wheat grain quality.

## Acknowledgments

The entire staff of Les Moulins de Soulanges and La Meunerie La Milanaise, and more specifically Élisabeth Vachon, Stéphanie Carrière, Chafik Baghdadi and Robert Beauchemin, are gratefully acknowledged for their support of this study. All members of the Labo Yergeau are thanked for their help in maintaining and setting up the field experiment. Emmy L’Espérance provided the R code used for repeated-measures ANOVAs.

